# Slow scission of single synaptic vesicles by Dynamin at physiological temperature

**DOI:** 10.1101/2025.03.18.643914

**Authors:** Sai Krishnan, Julia Lehrich, Yaroslav Tsytsyura, Nataliya Glyvuk, Junxiu Duan, Jürgen Klingauf

## Abstract

Compensatory endocytosis is crucial for the presynapse to maintain a functional pool of fusion-competent vesicles. Slow Clathrin-mediated endocytosis has been widely regarded as the primary mechanism of retrieval but this has been challenged by competing Clathrin-independent endocytic models, most notably sub-second ultra-fast endocytosis, reported to be predominant at physiological temperature. Here, we sought to resolve the salience of the respective endocytic modes by using a purely presynaptic preparation, the Xenapse, amenable to total internal reflection fluorescence microscopy (TIRFM). While the role of Clathrin is in dispute, Dynamin is widely acknowledged to figure as the scission protein at the invaginated vesicle neck. Hence, we labelled the endogenous Dynamin I with EGFP by CRISPR-Cas9 techniques and visualized single synaptic Dynamin-mediated scission events at very high temporal resolution. This revealed only a single slow mode of Dynamin-dependent retrieval with a half time of ∼ 9 seconds at physiological temperature. Cross-correlational analysis with fluorescently labelled Clathrin confirmed these Dynamin events to be Clathrin-dependent. We thereby affirm Clathrin-mediated endocytosis as the primary mode of compensatory retrieval.

## Main

In recent years, the role of Clathrin mediated endocytosis (CME) as the primary mode of compensatory endocytosis in the presynapse^1–4^ has been challenged by a highly specific Clathrin-independent endocytic mode, ultrafast endocytosis (UFE)^5–7^. This is purported to occur within 100 ms of a single action potential (AP) at the edge of the exocytic active zone and curiously only at physiological temperature, with CME predominating only at room temperature. UFE has been put forward as a rapid mechanism to maintain membrane homeostasis following exocytosis and the lack of any protein sorting mechanism entails going through an endosomal pathway where CME is required to form fully sorted synaptic vesicles (SVs). For all its differences with CME, Dynamin I (DynI) is also the scission protein for UFE where the long form splice variant of DynI in its preassembled form is needed for the rapid sub-second response^7–9^.

To disentangle the roles of CME and UFE in the presynapse we combined our previously characterized presynaptic preparation, Xenapse, amenable to total internal reflection fluorescence microscopy (TIRFM)^10^ with CRISPR-Cas9 gene editing to endogenously label DynI. This enabled the study of triggered compensatory endocytosis at an unprecedented spatial and temporal resolution at UFE conditions (i.e. 1 or 2 AP stimulus at 37°C). We find Dynamin-dependent fission to occur as discrete, quantal events characterized by stepwise assembly of recruited Dynamin followed by rapid removal after scission. Cross-correlation analysis with labelled Clathrin revealed both spontaneous and stimulus-dependent Dynamin scission events to be Clathrin-mediated thereby challenging the salience of the Dynamin-dependent and Clathrin-independent UFE mode. We identify Dynamin to be recruited to sites of preassembled Clathrin with the readily retrievable pool (RRetP) of Clathrin-coated structures (CCS) providing an element of speed advantage for the rapid, high fidelity retrieval of fusion-competent SVs in response to single AP. We further established a role for actin as force-generator during endocytic pit maturation enabling the rapid formation of SVs.

### Gene-edited Dyn1-EGFP allows imaging of single endocytic events

Initially we overexpressed DynI-EGFP in Xenapses^10^ and performed TIRFM at 37°C. While DynI-EGFP displayed stimulus-dependent recruitment (Extended Data Fig. 1a), resolving single events proved impossible due to a large rapidly exchanging membrane-associated ectopic fraction. We sought to overcome this problem by gene editing the endogenous DynI with the insertion of a fluorescent protein, an approach which has been successful in non-neuronal cell lines^11–13^. The CRISPR-Cas9 based pORANGE approach^14^, designed specifically for post-mitotic cells, was employed to knock-in EGFP at a splice invariant region of DynI (Fig. 1a). While the insertion site is adjacent to an intramolecular interaction interface, targeting this site was deemed safe as disruption of this interface was shown to result in only mild functional alteration^15^.

**Fig. 1:**
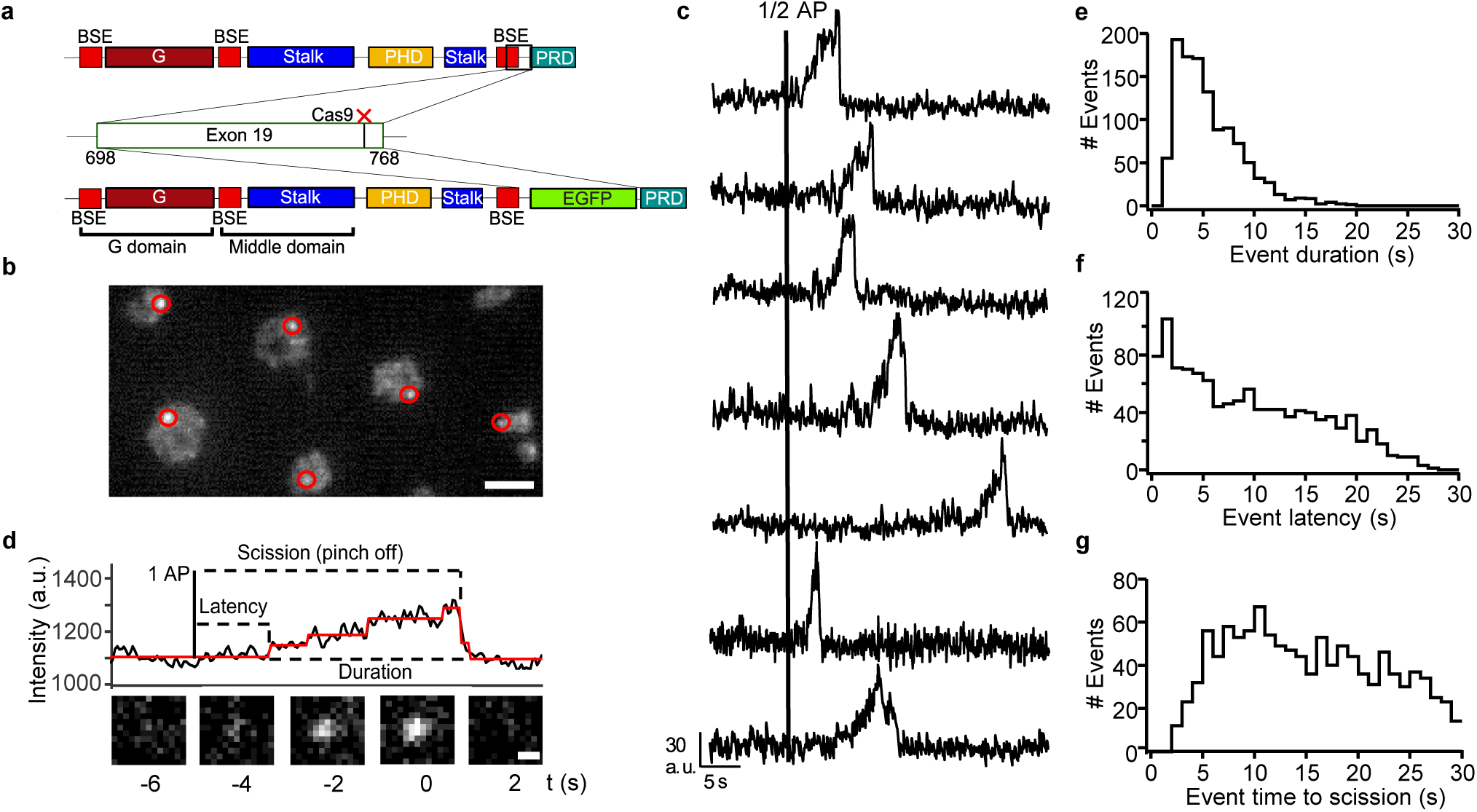
Recording single synaptic endocytic events with endogenously-tagged Dynamin I. **a,** Schematic depicting EGFP targetting to the Dynamin I gene by CRISPR-Cas9 based pORANGE strategy (BSE : Bundle signalling element, PRD : Proline rich domain). **b,** TIRF-microscopy image 3 s after 1 AP stimulus of Xenapses expressing EGFP tagged to endogenous Dynamin I with the Dynamin puncta circled in red; scale bar, 3 µm. **c,** Representative traces of individual evoked Dynamin recruitment and scission events occurring at various time points after the stimulus. **d,** An exemplar trace showing the stepwise Dynamin recruitment and annotated with measurable properties. Fit (red) obtained by applying the Autostepfinder step-detection function. Shown below are fluorescence images of the single Dynamin scission event at times indicated; scale bar, 0.5 µm. **e,** Event duration distribution, **f,** Event latency distribution and **g**, Event time to scission distribution. **e,f,g** are from the same data set of 1129 events in response to 1 or 2 AP and analyzed from n = 16 measurements from 7 biological replicates.

The endogenously expressed Dyn1-EGFP was recruited to the membrane in response to 1 or 2 APs in clear diffraction limited spots and was readily discernible due to the low background (Fig. 1b). Fluorescence time courses of single fission events display the distinct recruitment pattern also observed in non-neuronal single Dynamin fission studies^11,12,16^, a slow stepwise recruitment with a final spurt which is followed by sharp drop to the baseline (Fig. 1c). This can be interpreted as assembly of Dynamin units at the invaginating pit followed by scission (pinch off), consistent with GTP hydrolysis favoring disassembly of Dynamin oligomers at the membrane^17,18^. Applying a step-detector algorithm^19^ on these individual traces allowed for the extraction of the Dynamin event properties such as duration, latency and time to pinch off (Fig. 1d). While the median duration is 4.75 seconds (Fig. 1e), DynI recruitment for half the scission events occurs within 5 seconds after the AP (Fig. 1f). The ensuing time to pinch off occurs over a longer period with the peak in pinch off occurring at 10 seconds after the AP, consistent with the endocytic kinetics measured under similar conditions in hippocampal synapses using the fluorescent exo-endocytosis reporter pHluorin^20^. Post-stimulus recruitment is strikingly rapid occurring within a second of the AP and strongly suggests the recruitment to a pre-sorted and preassembled RRetP^21–23^. The average trace of these single post-stimulus events shows a rapid stimulus dependent increase followed by slower decrease of fluorescence (Extended Data Fig. 1b). While this trace shows bleaching, this is unlikely to be detrimental in determining the single event properties as most of these are only few seconds in duration. The average traces also do not show a drop in fluorescence after the stimulus, arguing against a role of preassembled DynI in compensatory endocytosis as has been claimed by a recent study^7^. Furthermore, the event duration also shows a distinct temperature dependency (Extended Data Fig. 1c) with a doubling of the duration to 9 seconds at room temperature which is consistent with Dynamin scission being enzymatically driven^24^.

### Dynamin mediated scission events in Xenapses are associated with Clathrin at physiological temperature

We next examined whether these Dynamin scission events represent CME by co-imaging of Dynamin and Halo-tagged neuronal Clathrin light chain (HaloTag-LCaI) at UFE conditions of 37°C and in response to 1 and 2 AP. In contrast to the clearly visible single Dynamin spots (Fig. 2a), Clathrin (Fig. 2b) events were difficult to discern as they are preassembled on the surface membrane at a high density, consistent with unroofed Xenapses revealing Clathrin pits decorating the inner membrane surface (Fig. 2c). This necessitated averaging of Clathrin events in space and time (time of fission) using the Dynamin signal as a reference, essentially performing cross-correlation with respect to the Dynamin scission frame. The resulting average traces for Dynamin and Clathrin (Fig. 2d,e) reveal Clathrin removal from the membrane at the same time point as Dynamin scission. As illustrated in the schematic (Fig. 2e) this corresponds to the preassembled Clathrin moving out of the evanescent filled following Dynamin-mediated scission. While this is strong evidence in favor of Clathrin involvement under UFE conditions, single event Clathrin dependency detection is prevented by the high background signal as is evident in the fluorescence images below Fig. 2d and e. Disrupting Clathrin assembly with the drug Pitstop-2^25^ significantly affects the Dynamin assembly and scission profile (Extended Data Fig. 2a) with a shallower Dynamin recruitment profile and marked disruption to the post-scission kinetics (Extended Data Fig. 2b). The number of bona fide scission events (as exemplified in Fig. 1c) is substantially reduced in the presence of Pitstop-2, with 55 vs 11 bona fide events for control vs treated culture conditions for the same Xenapses in response to 2 and 5AP.

**Fig. 2:**
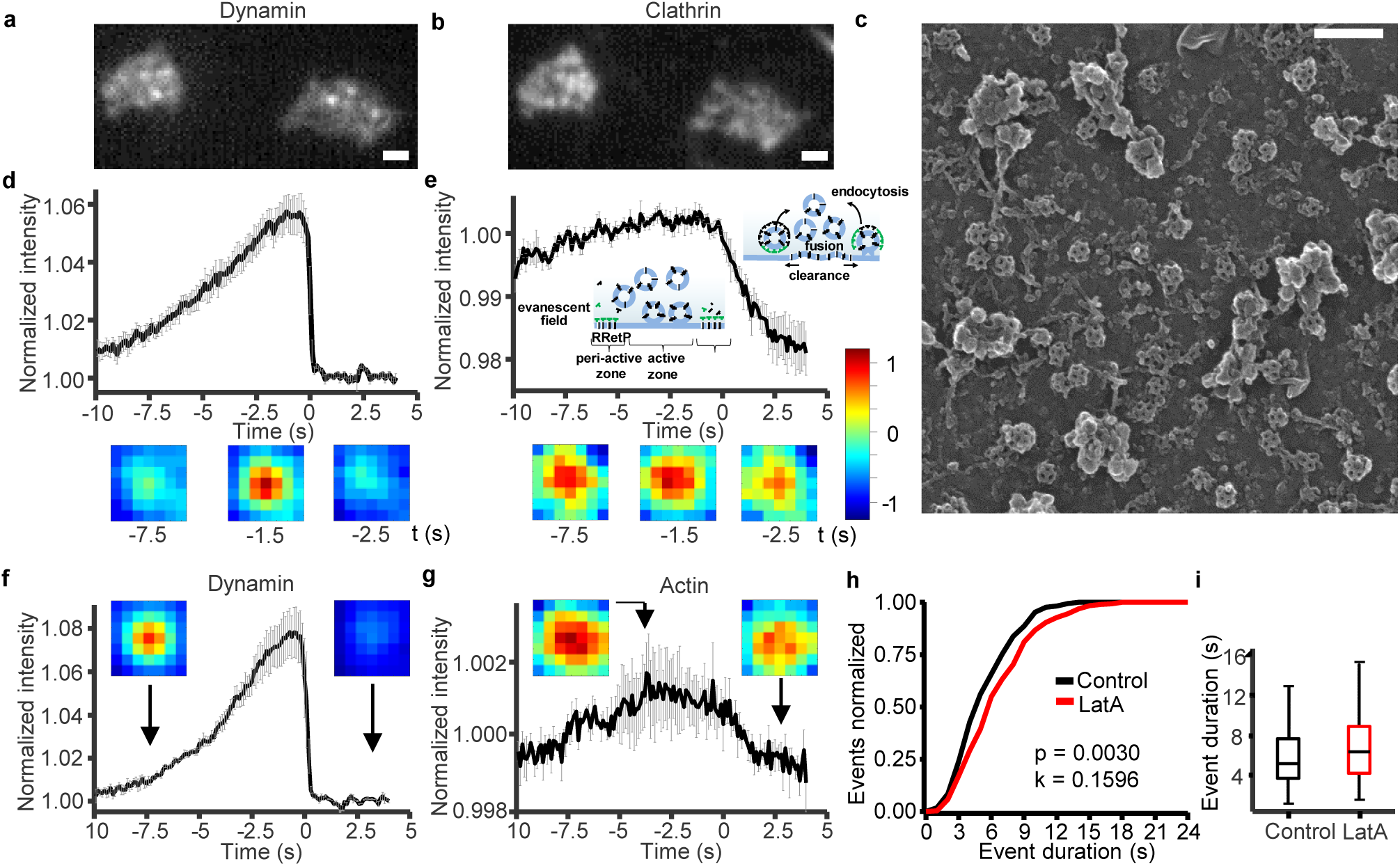
Dynamin events correlate with Clathrin removal and are sensitive to actin assembly perturbation. **a,b,** Representative fluorescence images (at 5 s after stimulus) of Xenapses co-expressing endogenously tagged Dynamin I-EGFP and overexpressed Halo-CLCa (stained with JF646) respectively; scale bar, 1 µm. **c,** SEM image of a section of an unroofed xenapse with its surface studded with Clathrin-coated structures. Scale bar 250 nm. **d**, **e**, Average trace of post-stimulus single Dynamin events centered in space and time along the scission frame **(d)** and of the corresponding average trace of Clathrin fluorescence **(e)**. Schematic in **(e)** shows the EGFP-labelled surface Clathrin within the evanescent field before and during endocytosis. Bottom: normalized fluorescence images of the averaged Dynamin event (**d**) and of the corresponding Clathrin events (**e**) at respective times. mean ± s.e.m. of 241 events (elicited at 1 or 2 AP) from n = 7 measurements and 3 biological replicates. **f, g,** Average trace of post-stimulus single Dynamin events centered in space and time along the scission frame (**f**) and corresponding average trace of tractin-mcherry fluorescence **(g)**. mean ± s.e.m. of 283 events (elicited at 2 or 5 AP) from n = 3 measurements and 2 biological replicates. Inset images show normalized average fluorescence at time points indicated by arrows. **h,** Normalized cumulative duration distributions of evoked events with LatA (red) and for control (black). 166 events for control from n = 2 independent measurements and for LatA 342 events measured in parallel from n = 6 measurements with both sets of data from 2 biological replicates with events evoked at 1 or 2 AP. **i,** Boxplot of the duration values in **h** with the central line showing the median and the bottom and top edges of the box indicating the 25th and 75th percentiles. The Kolmogorov-Smirnov test was used to show the LatA and No LatA control are from populations with significantly different distribution at the 1% significance level.

The role of actin in presynaptic fission has been insufficiently resolved^26–28^, unlike in non-neuronal cells where most studies agree about varying degrees of actin-dependency in promoting fission^12,26,29–31^. To address this we deployed the cross-correlation imaging approach by co-expressing of F-tractin-mCherry, a cellular marker for filamentous actin^32^. The average trace around the Dynamin scission frame for tractin showed noticeable decline preceded by accumulation of filamentous actin at the site of scission (Fig. 2f,g). Disruption of actin polymerization with LatrunculinA (LatA)^33,34^ displayed about 30 % prolongation of the Dynamin fission duration (Fig. 2h,i). This indicates that actin is indeed involved at the presynaptic scission site as an additional force generator during invagination and scission. This becomes clearer when looking at events averaged around the point of scission which shows the presence of LatA noticeably retards the time to scission following maximal assembly of Dynamin (Extended Data Fig. 3).

### Spontaneously occurring Dynamin scission events contribute to stimulus-dependent endocytosis

Xenapses like any synapse display constant low level baseline activity and we next sought to further analyze this pre-stimulus regime. This revealed spontaneous Dynamin events occurring entirely before stimulus (Fig. 3a) or ones where the assembly of Dynamin is initiated before but scission occurs following stimulus (Fig. 3b). The latter events form a special category of spontaneous events as they “straddle” the AP. Both spontaneous and straddle Dynamin events are Clathrin-mediated (Fig. 3c,d) and have similar duration distributions to the stimulus-induced events (Fig. 3e)underlining the ubiquity of Clathrin in mediating endocytosis presynaptically. When the waiting times to pinch-off (from stimulus) are plotted for both the post-stimulus and straddle events as a normalized inverse cumulative distribution (Fig. 3f), the time course of endocytosis is obtained. This distribution, after a four second lag, coinciding with the median duration of Dynamin buildup, decays monoexponentially with a time constant of about 10 seconds, indicating a single mode of retrieval. All this points to Dynamin and Clathrin mediated events occurring stochastically and they increase in frequency following a stimulus when there is an increased demand for recycling SVs.

**Fig. 3:**
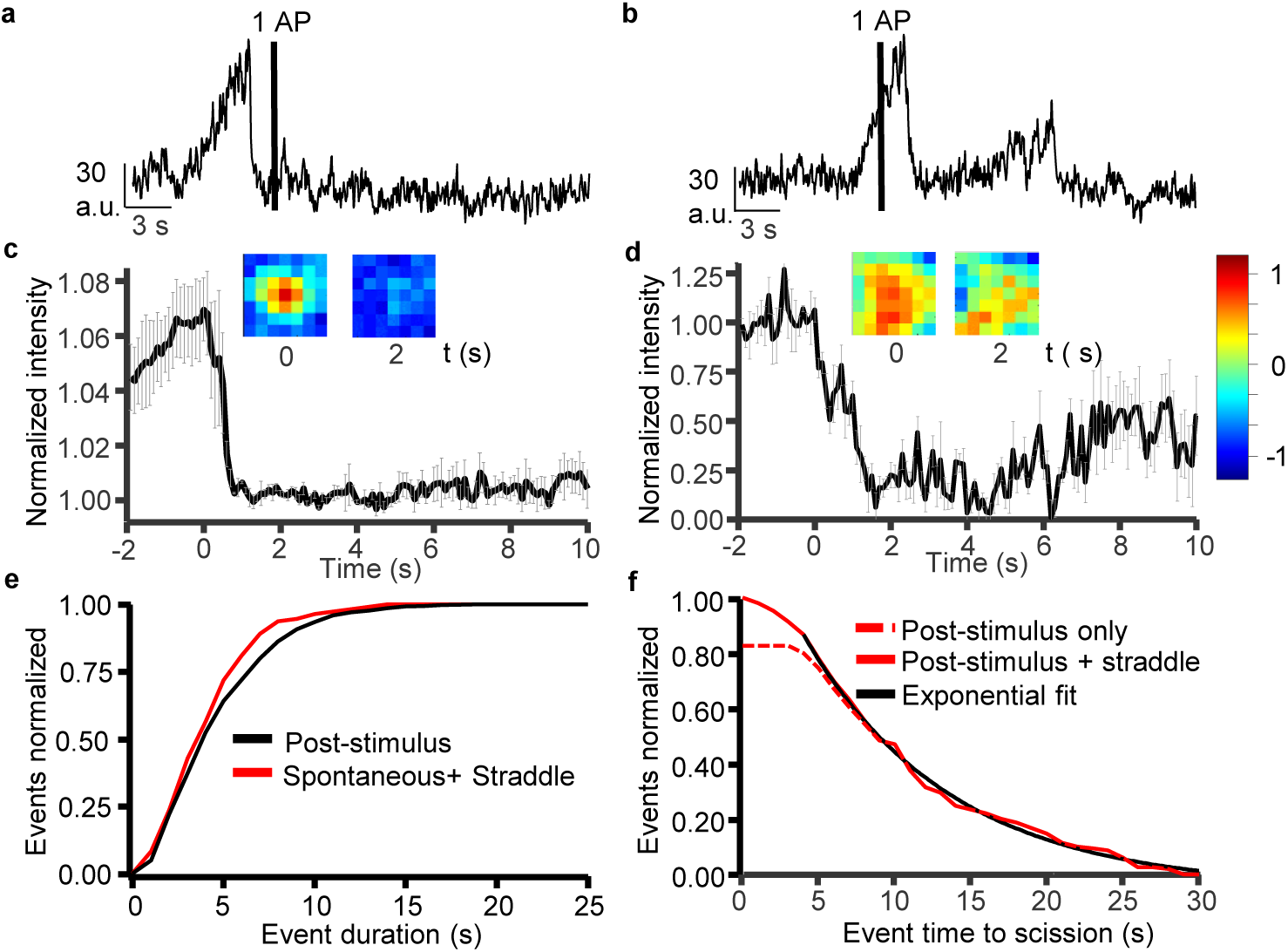
Spontaneous Dynamin events contribute to rapid stimulus-dependent endocytosis. Representative traces of pre-stimulus spontaneous **(a)** and straddle **(b)** Dynamin recruitment and scission events. **c, d,** Average trace of spontaneous single Dynamin events centered in space and time along the scission frame and the corresponding average trace of Clathrin fluorescence **(d)**. Inset images show normalized average fluorescence at time points indicated. mean ± s.e.m. of 38 events from n = 7 measurements with both sets of data from 3 biological replicates. **e,** Normalized cumulative event duration distributions for Dynamin I EGFP knock-in by pORANGE for evoked (black) and spontaneous events (red, including straddle events). 1031 evoked and 190 spontaneous events both from same data set consisting of n = 16 measurements and 9 biological replicates. **f,** Normalized inverse cumulative distribution of scission waiting times (time course of endocytosis) for all Dynamin scission events (solid red) and excluding straddle events (dashed red). A monoexponential fit (black) to the decaying phase after 4 seconds (median duration time) yields a time constant of 9.64 seconds. 206 evoked (1 AP) events and 27 straddle events both from n = 4 measurements and biological replicates.

The neuronally enriched DynI which is the focus of the study is well established to be differentially spliced at the C-terminus, resulting in splice forms DynaIXa and DynIXb^35,36^. While the C-terminal in the two splice forms differs by only 20 amino acids, this translates into functional differences with DynIXa being the splice form implicated in Clathrin-independent UFE^7,37^. We sought to test this by using the TKIT knock-in approach^38^ to target EGFP at the 3’end of the DynIXa to create a splice variant specific tagged Dynamin (Extended Data Fig. 4a). Comparing the TKIT tagged Dyn1Xa to the splice invariant pORANGE tagged DynI showed the same event behavior of slow assembly and scission (not shown) culminating in an almost identical duration distribution profile (Extended Data Fig. 4b). This suggests no functional specificity of DynIXa towards UFE, especially given nearly 55 % of the synaptic DynI labelled by pORANGE would be of the Xa form^35^. We further explored whether the large size of Xenapses formed on 5 µm spots would have any effect on Dynamin event dynamics by scaling the micropattern down to 2.5 µm spot size which results in Xenapse sizes (1.5 to 2 µm) comparable to large synaptic boutons found in normal hippocampal cultures (Extended Data Fig. 4c). Here too the event duration and the time to scission show no noticeable difference (Extended Data Fig. 4d,e) suggesting the size of the presynaptic bouton has no effect on the mechanism and kinetics of Dynamin recruitment and scission.

### Preferential recruitment of highly curved Clathrin coated pits revealed by latency distribution of Dynamin events

The first step increase of an event is taken to indicate the start of Dynamin incorporation at an endocytic site i.e. the latency (Fig. 1d). To understand the recruitment dynamics better we plotted the latency distribution, i.e. the time from stimulus to the first recruitment step, for Dynamin in response to 1, 2 and 5 APs. The latency distributions for 1 and 2 APs were similar. Both however show a slight shoulder at around 10 seconds. which becomes visible as pronounced hump for 5 APs indicating a second recruitment phase or two RRetP subpools from which endocytosing vesicles are drawn (Fig. 4a, average trace of events in Extended Data Fig.5). While for 1 AP most vesicles would be drawn from an immediately available RRetP, for 2 APs about half and for 5 APs the majority of vesicles would be drawn from CCS that are not fully assembled yet and would need about 10 seconds for completion before Dynamin can be recruited. This interpretation is supported by the structural distribution of the endocytic CCSs found on cortices from unroofed Xenapses. The CCS can be broadly classified based on their curvature state as domes and pits representing different stages of endocytic maturity (Fig. 4b). Dynamin is preferentially recruited to the higher curvature found at the invaginating neck^39^ and it is conceivable that Clathrin-mediated Dynamin scission occurs first at the highly curved pits consistent with the sub-second post-stimulus recruitment of Dynamin (Fig. 1g). At 5 AP when there is a greater endocytic demand, the less mature domed structures are activated for invagination but evidently it takes up to 10 seconds for it to achieve sufficient curvature to recruit DynI molecules. While this correlation of function to structure is indirect it is lent further credence by the observed time constant of surface Clathrin reorganization in response to FRAP also amounts to ∼10 seconds^40,41^. The significantly greater number of domed in comparison to pits further highlights the former Clathrin structures to be a potential back up during increased vesicular demand (Fig. 4c).

**Fig. 4:**
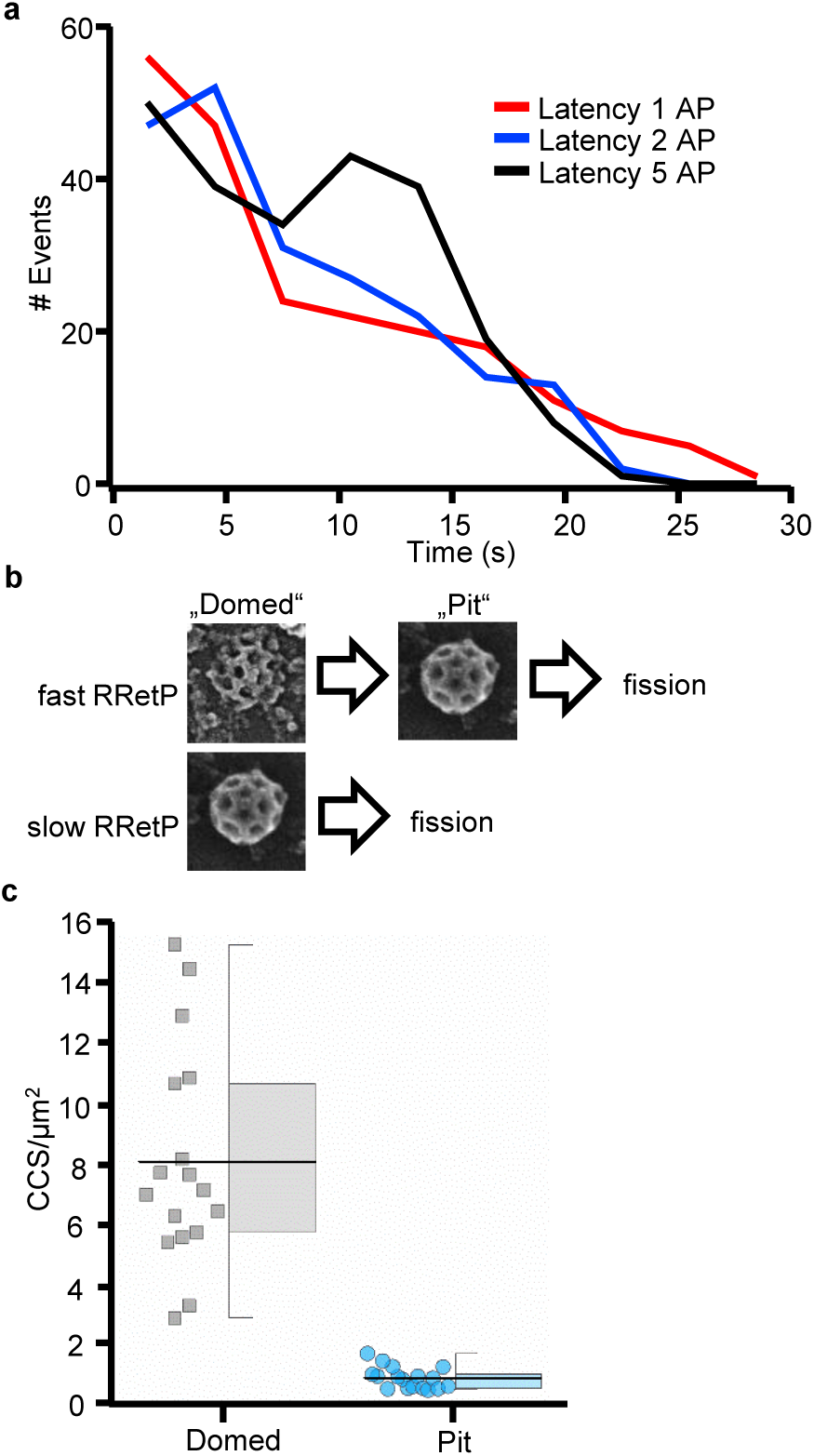
Latencies of Dynamin events at increasing stimuli reveal existence of slow and fast RRetP sub-pools and their structural correlates. **a,** Event latency distribution for Dynamin I-EGFP (pORANGE knock-in) scission events in response to 1 (n = 206), 2 (n = 208) and 5 AP (n = 233). n = 4 measurement from 3 biological replicates for 1 AP and n = 6 measurements from 5 biological replicates for 2 and 5 AP. **b**, SEM images of the two major observed classes of Clathrin coated structures, domed and pit, and the transition between the two. **c**, Density of domed (8.03 ± 3.62 per mm^2^) and pit (0.81 ± 0.36 per mm^2^) structures from SEM images of unroofed Xenapses (n = 17).

### Stoichiometry of Dynamin molecules at the site scission

Targeting a splice invariant region of DynI gave us the opportunity to measure the number of molecules that are incorporated into the vesicle neck at the point of scission. The fluorescence step size a single EGFP molecule was determined (Extended Data Fig. 6a) to enable calibration to the amplitude of single Dynamin events (Extended Data Fig. 6b). The intensity distribution of the single EGFPs gave a mean value of ∼16 a.u. with the mean value of Dynamin scission amplitude amounting to 160 a.u. This would indicate an average of 10 DynI molecules but performing gene editing in a post-mitotic culture does not allow for the selection of biallelic insertions and the single intensity distribution of DynI indicates monallelic insertion of EGFP. We can therefore assume an average of 20 DynI molecules is incorporated into the scission site and this could be extended to ∼25 molecules when we observe maximum monoallelic intensities greater than 200 a.u. This approximates the observed minimum number of dynamin per turn at the membrane neck, as demonstrated in Cryo-EM reconstruction images and live studies^11,12,42,43^. The estimated numbers could be an underestimate due to incorporation of the unlabeled DynIII isoform also found at the presynapse but at lower levels than DynI^44^. It also cannot be ruled out that a portion of the dynamin scission events with a lower amplitude fail to complete successful scission.

## Discussion

Here we provide direct evidence challenging the apparent role of UFE as the primary mode of membrane retrieval at the pre-synapse. Combining gene-edited DynI with overexpressed Halo-LCaI (stained with JF646) revealed single Dynamin events to be Clathrin mediated, thereby establishing CME as the primary mode of ensuring rapid and well-choreographed retrieval of the RRetP for the eventual formation of fusion-competent SVs. Our data showing CCS of varying size and curvature are in line with a ‘mixed’ model between constant area and constant curvature with a small subpool of complete CCP but also shallow domed structures that need to recruit more Clathrin and other accessory proteins but are endocytosed even after single AP^45–47^. The recruitment of Dynamin to the preassembled CCP suggests assembly of Dynamin at the invaginating vesicle neck could be the rate-limiting step during endocytosis of single SVs.

The direct visualization of Clathrin mediated scission under UFE condition is in contrast to the evidence adduced in favor of UFE which has relied almost exclusively on electronic microscopy images done on neuronal sections subjected to optogenetic flash-freeze. A peculiar feature of UFE invaginations is their volume is typically 4-times larger than that of a single SV, despite occurring in response to 1-2 APs. Rapid endocytosis of uncoated vesicles with four times the area of SVs at the edge of the AZ was observed in frog neuromuscular junction (NMJ) already 40 years ago^48^. About a third of the exocytosed membrane was retrieved by this fast mechanism, which the authors attributed to an unphysiological excess amount of SV membrane added to the plasma membrane in the presence of the K^+^ channel blocker 4-AP, that was used to prolong the single AP applied prior to plunge freezing. It is therefore possible that the UFE invaginations could be the result of Channelrhodopsin eliciting prolonged neuronal depolarization.

Freeze-fracture replicas of frog NMJ revealed that these enlarged structures, unlike small coated pits, have no capacity to enrich large particles^48^. The authors noted that this process may thus explain the appearance of “cisternae”, endosome–like intermediates, after an unphysiological train of stimuli. Indeed, the endocytosed vesicular structures in UFE have to proceed via an endosomal pathway to achieve proper cargo sorting by CME and generate fusion-competent SVs^6^. This undermines the apparent speed advantage of UFE but the authors justify this by claiming rapid retrieval to be necessary to rapidly reset membrane area. A well-established Clathrin-independent mode such as activity dependent bulk endocytosis (ADBE) also passes through an endosomal intermediate but is perfectly compensatory in membrane retrieval^49–51^. In this case the need for rapid membrane retrieval is more obvious as ADBE is elicited by high K^+^ or high frequency stimulation. The lack of cargo-sorting of UFE however would lead to an imbalance of membrane lipid and SV protein recycling, leaving cargo to accumulate at the surface. This imbalance in membrane lipid and protein trafficking theoretically can only be resolved by an additional retrieval mechanism with an even higher sorting capacity than CME working in parallel and/or a high fusion rate of ‘empty’ endosomal intermediates, from which SV proteins had been removed by CME. Neither process has been described thus far. The molecular mechanism put forward to enable the rapid uptake in UFE is that the DynI may be preassembled at the periactive zone which would allow it to instantly assemble at the invaginated vesicle neck^7^. The presence of puncta at the membrane surface is presented as evidence for preassembly but we observe these only for DynI overexpression as ectopic highly dynamic structures which do not respond to stimuli. In any case, our observation that spontaneous Dynamin scission contributes to the post-stimulus endocytic response (Fig.3f) makes the need for preassembly redundant.

In challenging UFE, we have presented here the first direct visualization of single SV endocytosis events and their underlying properties. While the recruitment and scission behavior of the events was broadly similar to those in non-neuronal cells, synaptic scission was noticeably different in its duration being shorter and having a narrower distribution^11,12^. This was reflected in the stoichiometry of Dynamin molecules which at most form a little more than a turn at the vesicle neck (Extended data Fig.6) while Dynamin can assemble to up to 3 turns in non-neuronal fission sites^12^. All this is consistent with the need to form SVs of uniform morphology with defined quantal content and points to possible role for dynamin in shaping these regularly sized SVs. The substantial disruption to post-scission Dynamin decay following treatment with Pitstop-2 (Extended data Fig.2) suggests Clathrin coat assembly itself to be a contributor to the invaginating force during endocytosis, consistent with studies showing Clathrin to drive spontaneous membrane curvature^52–54^.

The sculpting role in scission also likely involves accessory proteins such as actin, BAR domain proteins such as Amphiphysin I and ANTH domain containing Epsin I, proteins which are known to directly or indirectly influence Dynamin function^55–58^. Our use of tractin-mCherry shows actin assembly during scission to precede maximal Dynamin assembly, consistent with non-neuronal observations which show actin by arriving earlier to the scission site promotes Dynamin recruitment. The mechanism likely involves actin promoting invagination by acting as a force generator with a potential role for a feed-back loop mechanism indirectly promoting Dynamin recruitment. This role as force generator seems to continue during the scission process as reflected in the increased duration for initiation of scission once Dynamin is fully assembled (Extended data Fig. 3). The *in vivo* dynamics of BAR domain proteins and Epsin I in the presynapse remains unknown. Our novel approach of studying triggered single synaptic scission events at high precision opens the prospect of investigating the choreography of other key endocytic proteins relative to Dynamin fission.

## Methods

### Animals

Breeding of C57BL/6N mice was conducted at the Central Animal Experimental Facility of the University Hospital Muenster according to the German Animal Welfare guidelines (permit number AZ81–02.05.50.20.019, issued by ‘Landesamt für Natur, Umwelt und Verbraucherschutz NRW’, Duesseldorf, Germany). Both male and female mice were used for primary neuronal cultures.

### Plasmid constructs

Dynamin1-EGFP overexpression plasmid was provided courtesy of Prof. V. Haucke. Halo-CLCa was derived from EGFP-CLCa construct with its neuron specific Clathrin light chain (LCaI) which has been described previously^23^. Restriction digestion removal of EGFP from the latter construct was followed by insertion of Halo from PCR amplification of Neuroligin1ΔB-Halotag-IgG construct provided by the lab of Prof. M. Missler. Tractin-mCherry plasmid was a gift from the lab of Prof.Tobias Meyer (Addgene plasmid # 85131).

Dynamin I in primary neuronal cultures was gene edited to insert EGFP using both the pORANGE and TKIT approaches^14,38^. The 20 bp gRNA duplexes were generated by stepwise annealing of the forward and reverse sequence with a temperature gradient of 5°C/min from 95°C to 25°C. These were then individually ligated into BbsI digested pORANGE cloning template vector (Addgene Plasmid #131471) enabled by the duplex having CACCG and AAAC overhangs at either end. The single exonic gRNA duplex for pORANGE and the two intronic gRNA duplexes for TKIT targeted distinct loci of the endogenous Dynamin I (Fig.1A and Fig.3G). The TKIT gRNAs were combined in a single vector by PCR amplification of one of the gRNAs along with its flanking U6 promoter and gRNA scaffold and its insertion at the pORANGE XbaI site, 3’ to the other gRNA expression cassette.

The EGFP donor for pORANGE was generated by PCR amplification with primers flanked by reverse complement of the genomic target sequences and was inserted into the pORANGE vector at the HindIII and XhoI site. For TKIT, a donor containing the exon and in frame EGFP flanked by the intronic sequences and the reverse complement of the genomic target sequences was generated as a synthetic DNA fragment (Eurofins Genomics GmbH, Ebersberg). This was subsequently inserted into the HindIII and XhoI site of the pORANGE vector containing the two gRNA fragments. The lentiviral plasmid for pORANGE was generated by PCR amplifying the knock-in cassette containing the U6 promoter, gRNA duplex, and donor DNA. This was inserted into PacI digested pFUGW vector (pFUGW mCherry-KASH, Addgene Plasmid #131505) using Gibson cloning (NEBuilder HiFi DNA Assembly).

All primers used in cloning the above constructs are listed in the Supplementary Table 1. Sequence of all cloned plasmids were confirmed by Sanger sequencing at Microsynth Seqlab GmbH, Göttingen.

### Transfection and Transduction

Constructs purified by Maxi-prep (Qiagen, Germany) were used to transfect neurons at 4 days *in vitro* using a modified version of previously described Ca^2+^ Phosphate transfection protocol^59^.

Lentiviral particles for pORANGE Dynamin1-EGFP knock-in were generated in HEK cells by combining the gRNA cassette containing pFUGW mCherry-KASH with the lentiviral envelope (pMD2.G) and packaging (pCMVΔR8.2) plasmids which were gifts from the lab of Prof.Didier Trono (Addgene plasmids # 12263 and #22036 respectively). Lentiviral viral particles were produced as per our established protocol^59^. Neurons were subjected to 6 hour incubation in fresh growth media with the viral particles in a 5% CO_2_ incubator at 37°C. Following HBSS wash, the coverslips were transferred to the original plate.

### Cell culture and imaging conditions

Xenapses were generated from dissociated hippocampal neurons from newborn (P0) C57/BL6 mice cultured on 18 mm micropatterned glass coverslips functionalized with Neuroligin 1(+A ΔB) and placed in a Banker-type culture^10^. Briefly, PDMS stamps were generated with customized micropatterned surface corresponding to the desired Xenapse dimensions (5 µm diameter, 8 µm center to center distance or 2.5 µm diameter, 5.5 µm center to center distance).

The stamps were inked with PLL-PEG-HTL (PLL grafted with PEG-HTL) and transferred onto plasma cleaned glass coverslip surface. The resulting coverslip was micropatterned with the PLL-PEG-HTL and readily bound the Halotag containing Neuroligin I, which in turn could bind its endogenous partner Neurexin^60^ in hippocampal neurons thereby resulting in the generation of presynaptic Xenapse culture.

All imaging was carried out at 14-18 days *in vitro* with cells placed in measuring buffer (140 mM NaCl, 2.4 mM KCL, 4 mM CaCl_2_, 1.3 mM MgCl_2_, 10 mM HEPES, and 10 mM D-Glucose, pH adjusted to 7.4) containing 50 µM D, L-2-amino-5-phosphonovaleric acid (AP5), and 10 µM 6-cyano-7-nitroquinoxaline-2, 3-dione (CNQX) to prevent recurring neuronal activity. The elevated Ca^2+^ concentration was used to mimic the UFE measuring conditions^5^. Actin assembly was disrupted by addition of 50 µM LatrunculinA (Tocris Bioscience) in measuring buffer followed by 5 minute incubation before commencing image acquisition. Clathrin assembly was disrupted by addition of 30 µM Pitstop-2 in measuring buffer and acquisition initiated within a minute of addition. All imaging used 10 ms exposure and acquisition was at 100 Hz for non-optosplit measurements and 50 Hz for optosplit measurements due to toggling between the two laser channels. The overexpressed Dynamin1 measurement (Extended Data Fig.1) was acquired at 1 Hz with a 50 ms exposure.

### TIRF microscopy and fluorescence calibration

TIRF illumination was achieved using a Nikon Eclipse Ti microscope and 100x / 1.49 NA Apo TIRF oil immersion objective. A cooled scMOS PRIME 95B camera (11 µm x 11 µm pixel area, Photometrics) was side mounted onto the microscope and image acquisition controlled by a Micromanager imaging software (https://micro-manager.org/).

Coverslips were mounted on chamber with platinum electrodes to achieve electric field stimulation by the application of 80 mA pulse administered by a constant current stimulus isolator (WPI A 385, World Precision Instruments). OMICRON SOLE-6 laser light engine with high-speed analog and digital modulation controlled the lasers. The relevant fluorescent constructs used in this study were illuminated with either 488 nm, 638 nm (both diode lasers) or 561 nm (diode-pumped solid-state laser).

Dual colour simultaneous imaging was achieved by a filter cube inserted into an Optosplit II image splitter (Cairn Research, UK) with its output port fitted to the camera. The optical path of the filter cube had inserted in it a 583 nm dichroic and 685/70 nm and 525/45 nm bandpass filters (AHF analysentechnik AG, Tübingen, Germany) to allow for the separation of the GFP and JF646 fluorophore emission. The aperture screws in the image splitter were used to form two roughly equally sized rectangular images, from the two halves of the camera sensor, allowing for the acquisition of the fluorophores without bleed through.

Measurements at 37°C was performed by placing the measuring chamber in the Heating insert P and Incubator PM 2000 on the microscope mechanical stage, which in turn were connected to the Tempcontrol 37-2 Digital Microscope Temperature Process Controller (PeCon Gmbh).

An Arduino microcontroller unit (Arduino IDE 1.6.0) with a self-written code functioned as the external trigger controlling with high precision the lasers, camera, and stimulus isolator.

### Image analysis

All detection of Dynamin events and extraction of data was performed using the 64-bit version of Matlab 2019b (MathWorks, Natick, MA, United States of America). The first step in detecting single Dynamin events was background subtraction following image read-in. This was achieved by averaging the pre-stimulus frames, or the post stimulus frames for spontaneous events and subtracting them from the rest of the movie. Peak fitting was done with a 3×3 pixel region on the background subtracted movie found by *’à trous*’ wavelet filtering ^61^ using a pre-determined PSF template with subpixel accuracy as described previously^10^. The localizations were then filtered using an intensity-based mask to include synaptic boutons only and by removing overlapping localizations within five pixels in the same frame. Then localizations within the same 3×3 pixel area were filtered in time by selecting the only brightest events in order to remove duplicates. The resulting localizations were applied on the original movie to generate intensity traces of single Dynamin events.

The individual median-filtered traces were subjected to an automated step-fitting procedure using a publicly available step-fit algorithm, AutoStepfinder^19^. This used an iterative process which minimized the variance between data and fit, based on the assumption that the data contains instantaneous steps. The user-interface allowed the setting of a threshold to define the quality of the fit which was typically determined by the length of the trace being analyzed. The step properties were automatically saved as a Matlab file and could be further processed to extract the relevant properties which were the frames at which the steps of an event occur and their respective sizes recorded in arbitrary fluorescence intensity units. The first and last frame determined the latency and scission respectively relative to the stimulus frame and the difference between the two gave the duration of the event.

For the stoichiometry calculation, coverslips cleaned with Piranha solution (10 minutes incubation in concentrated sulfuric acid with hydrogen peroxide in a ratio of 3:1) was plasma cleaned and functionalized as described above with the exception of EGFP-Halo (kindly provided by Prof.Jacob Piehler of Osnabrück University) instead of Neuroligin I-Halo was immobilized at a low surface density. Single step EGFP bleaching events were recorded at the same laser and acquisition settings as the Dynamin measurements. This measurement was done in parallel to the Dynamin measurement to minimize any drift in laser intensity. The Dynamin events acquisition itself was performed at lower duty cycle (10 ms exposure at 10 Hz acquisition) to reduce bleaching and thereby determine a more accurate estimate of events amplitude i.e the final scission step size. The single EGFP molecules were closer to the coverslip surface relative to the Dynamin-EGFP in Xenapses by at least 21 nm (synaptic cleft width of about 15 nm, defined by the Neuroligin-Neurexin complex, and the plasma membrane thickness of 6 nm). An intensity correction was therefore performed by reducing the recorded fluorescence units of single EGFP step sizes by 20 % for an evanescent field depth of 100 nm ^62^. The histogram distribution of the corrected step sizes of the single EGFP relative to the size distribution of the final scission step of Dynamin events was used to determine the stoichiometry of Dynamin molecules involved in scission.

The correction for shift between the two rectangular halves representing the two channels of the optoplit measurement was achieved by a self-written Matlab script. Images of 0.5 µm broad spectrum Tetraspeck^TM^ beads (Invitrogen), deposited on plasma cleaned coverslips at low density, and were taken in parallel with the optoplit movie acquisition and the two equally separated halves of the bead images were read-in for correction. The resulting coordinate shift was applied on the right channel of the Dynamin measurement to correct for the shift relative to the left channel.

### Electron microscopy of unroofed xenapses

Unroofing of Xenapses at DIV 16-18 was performed according to established protocols^63,64^. In brief, excess media was washed off the cells with calcium-free HEPES-buffered Ringer solution (in mM: NaCl 140, KCl 2.4, MgCl_2_ 4, HEPES 10, D-glucose 10, pH 7.3), then incubated briefly in the same solution containing 0.1 mg/ml PLL. After washing off the excess PLL, cells were placed in hypotonic PHEM buffer (in mM: PIPES 60, HEPES 25, MgSO_4_, EGTA 10, pH 6.96/KOH; 1 part of buffer mixed with 2 parts of deionized water). Unroofing was carried out in PHEM buffer using Branson 250 ultrasonifier with a 13 mm tip attached to the resonator at the lowest strength for a sub-second pulse duration.

Unroofed cells were instantly fixed in PHEM buffer containing 1-2% glutaraldehyde (GA) (TEM grade, Serva GmbH, Germany) for 15 minutes at room temeprature. Samples were fixed in aqueous tannic acid 0.2% (Sigma Aldrich, USA) and stabilized with aqueous FeCl_3_ 0.2% for 15 minutes each with extensive washing with deionized water preceding and following each step. Cells were dehydrated in ethanol series (50%, 70%, 2x 100%, 1 minute each, agitated), briefly incubated in 1:1 mixture of 100% ethanol with hexamethyldisilazane (HMDS, Carl Roth, Germany), transferred to pure HMDS (twice, 1 min each, agitated) and then air dried. Unroofed Xenapses were identified using phase-contrast light microscopy, isolated with a diamond cutter, glued on the Al sample carriers with CCC (‘Conductive Carbon Cement after Göcke’, Plano GmbH, Germany), and rotary coated after full drying of cement with Pt/C 2 nm at 45° in Leica ACE900. Scanning electron microscopy was performed with FEG SEM Hitachi S5000 (Hitachi, Japan) at 30 kV using a secondary electron detector; images were acquired and digitized using DISS5 (Point Electronic, Germany). The data analysis (CCP size and type distribution) was performed in MetaMorph (Molecular Devices, USA) and OriginLab (OriginLab Co, USA).

Supplementary Information is available for this paper.

**<u>Supplementary Table 1.</u>** List of primers used for generating the constructs used in this study.

## Acknowledgements

We thank K. Tkotz and A. Roetrige for excellent technical support. We also thank M.Kahms for proofreading the manuscript. This work was supported by grants of the Deutsche Forschungsgemeinschaft (SFB 944 and SFB 1348) to J.K.

## Author information

The authors declare no competing financial interests. Correspondance and requests for materials should be addressed to J.K.

## Author contributions

S.K. and J.K. conceptualized the study and designed the experiments; S.K. conducted the imaging experiments supported by J.D; S.K. and J.K. analyzed the imaging data; N.G and Y.T conducted and analyzed EM experiments, J.L. wrote software for data analysis; S.K. and J.K. wrote the manuscript.

## Data availability

The data that support the findings of this study are available from the corresponding author upon reasonable request.

## Code availability

Matlab Code used for analyzing the data presented in this manuscript can be accessed from GitHub with the following DOI: 10.5281/zenodo.13934548.

## Figure Legends

See pdf of figures

---

**Extended Data Fig.1:**
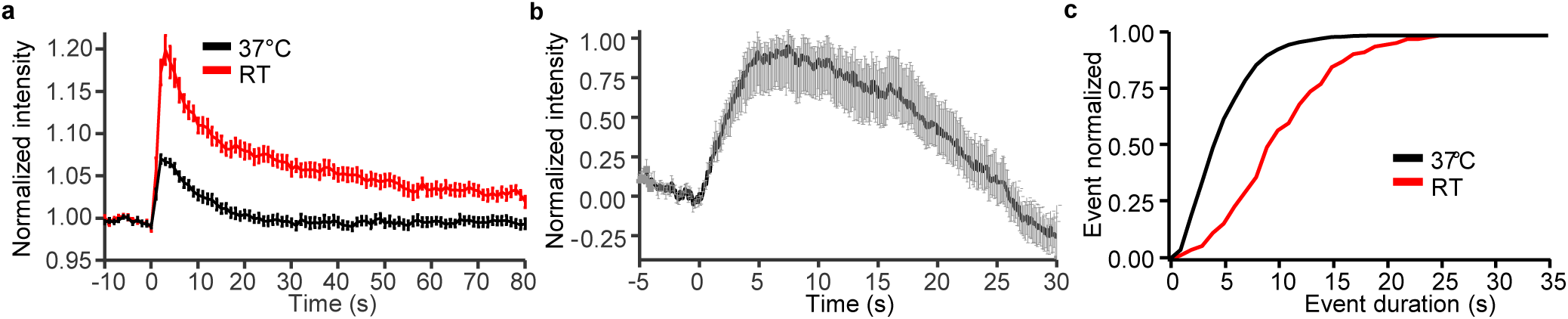
Temperature dependency of Dynamin I scission dynamics. **a,** Normalized average fluorescent traces of Dynamin I-EGFP overexpressed in Xenapses responding to a 50 AP 40 Hz stimulus at RT (black) and 37°C (red). mean ± s.e.m. of n = 19 measurements at RT vs n = 27 measurements at 37°C both from 5 biological replicates. **b,** Normalized average trace of single Dynamin events elicited with 1 or 2 AP. mean ± s.e.m. **c,** Normalized cumulative event duration distribution (pORANGE Dynamin I-EGFP knock-in) for RT (red) and 37°C (black). For 37°C, 1031 events (evoked with 1 or 2 AP) consisting of n = 16 measurements from 9 biological replicates. For RT, 119 events (evoked with 2 or 5 AP) consisting of n = 10 measurements from 2 biological replicates.

**Extended Data Fig.2:**
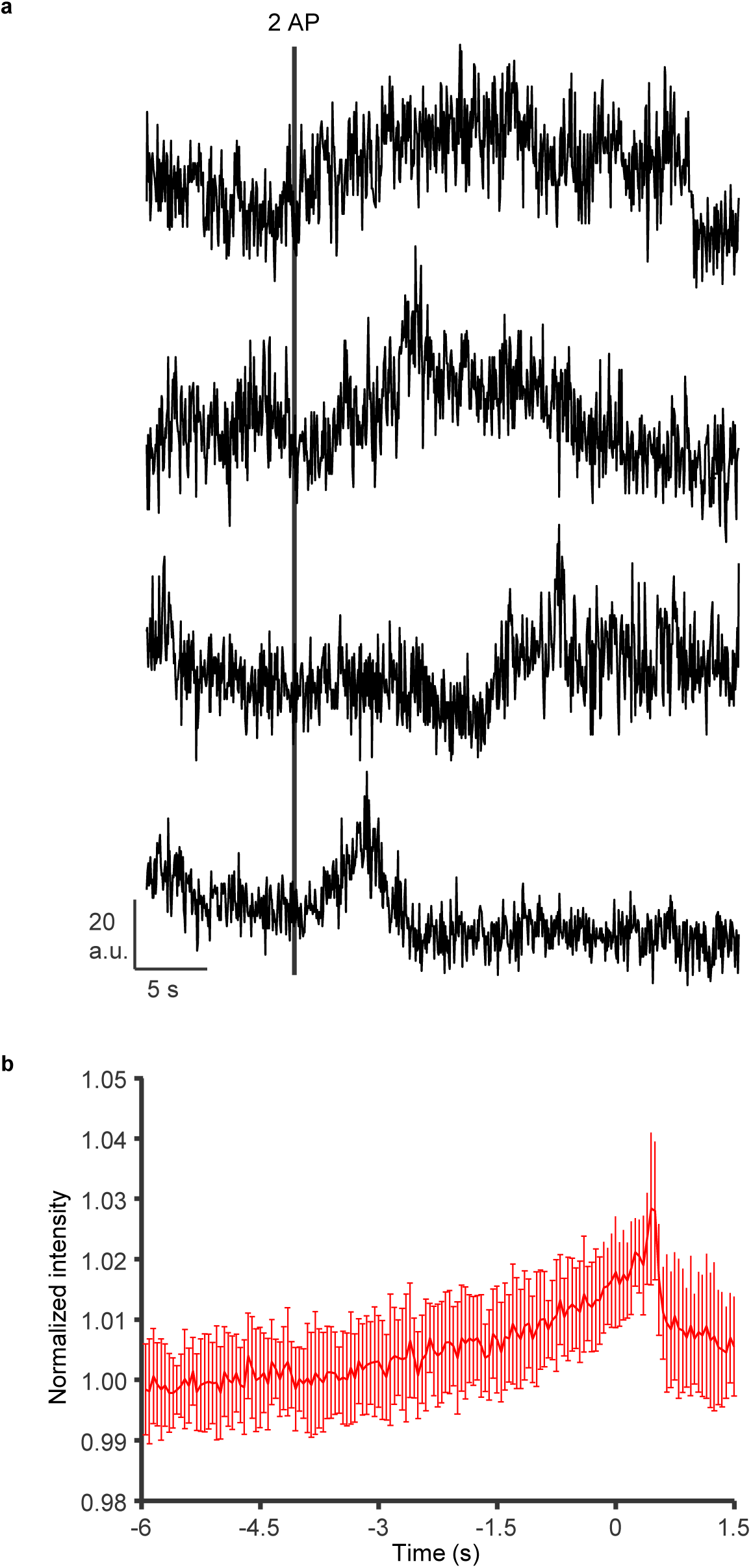
Disruption of Dynamin I scission dynamics by Pitstop-2 mediated inhibition of Clathrin assembly. **a,** Exemplary traces of single Dynamin events in the presence of Pitstop-2 (30 µM) with their assembly or scission profiles disrupted. **b,** Average trace of post-stimulus single Dynamin events centered in space and time along the scission frame for Pitstop-2 (30 µM) treated events. 51 events from n = 4 measurements and 2 biological replicates with post-stimulus events evoked at 2 and 5 AP.

**Extended Data Fig.3:**
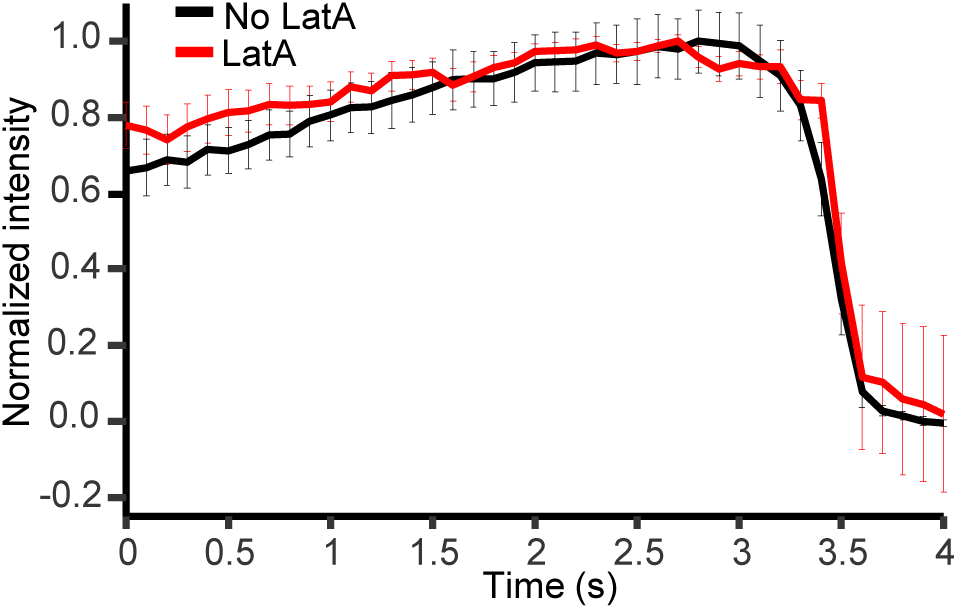
Disruption of Dynamin I scission dynamics by LatA mediated inhibition of actin assembly. Average traces of post-stimulus single Dynamin events centered in space and time along the scission frame from the data set in **Fig.2h** with (red) and without (black) LatA treatment.

**Extended Data Fig.4:**
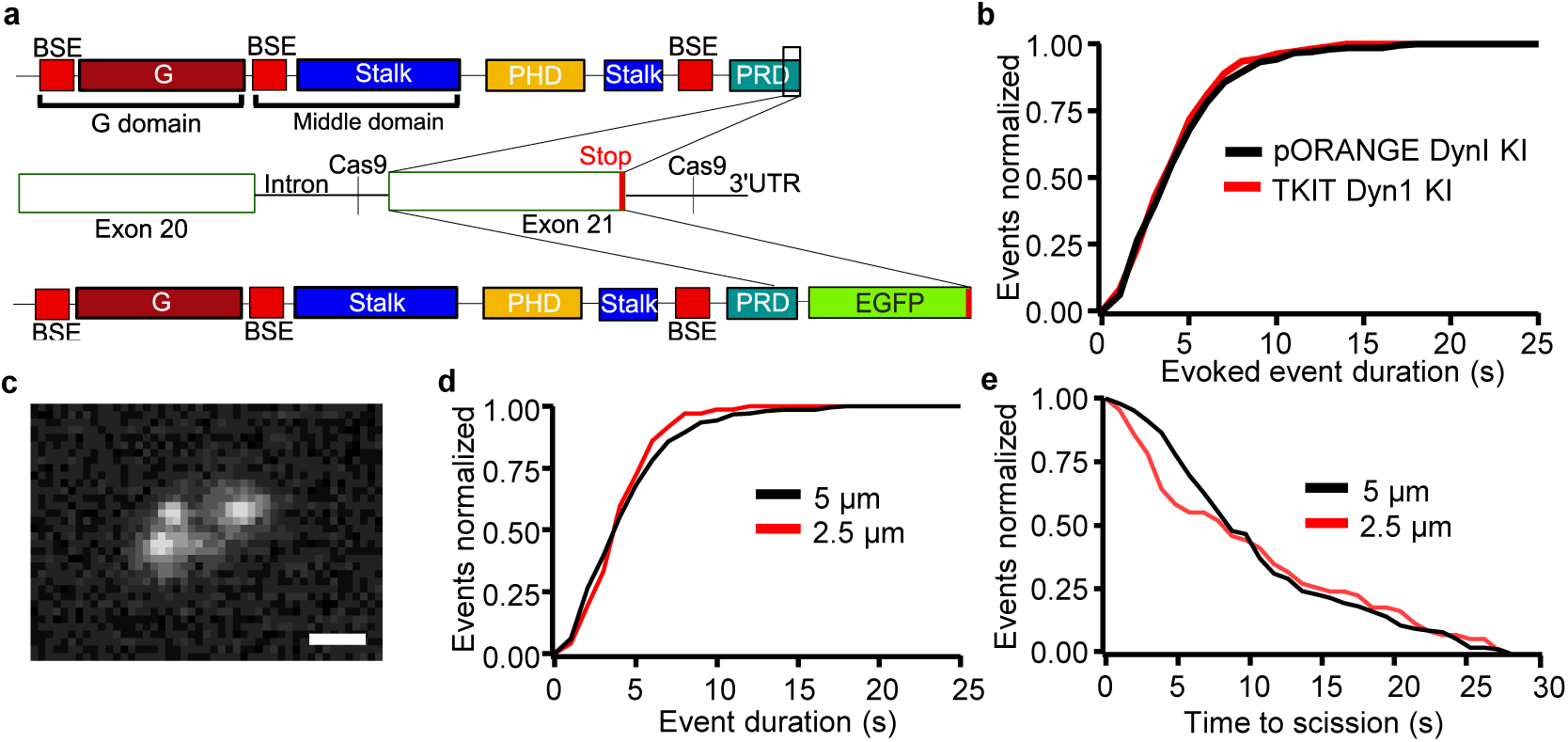
Dynamin event properties conserved across splice variants and bouton size. **a,** Schematic depicting EGFP insertion to the Dynamin I 3‘UTR by CRISPR-Cas9 based TKIT strategy. **b,** Normalized cumulative stimulus-evoked event duration for Dynamin I EGFP-tagged by pORANGE (black) and TKIT(red) knock-in strategies. 118 events from n = 6 measurements and 2 biological replicates with post-stimulus events evoked at 2 or 5 AP.**c,** Representative TIRFM image of a Xenapse formed on a 2.5 µm micropattern expressing pORANGE tagged Dynamin-EGFP. Several puncta of Dynamin recruitment are visible 1 s after 2 AP; scale bar, 1 µm. **d,** Normalized cumulative event durations including straddle events for Xenapses growing on 5 µm (black) and 2.5 µm (red) micropatterns. **e**, Normalized inverse cumulative distribution of waiting times of scission (time course of endocytosis) for Xenapses growing on 5 µm (black) and 2.5 µm (red) micropatterns. For pORANGE **(b)** and 5 µm spots **(d,e)**, same data set as for post-stimulus in **Fig.3f**. and for 2.5 µm spots **(d,e)**, 65 events from n = 7 measurements and 2 biological replicates with post-stimulus events evoked at 1, 2 or 5 AP.

**Extended Data Fig.5:**
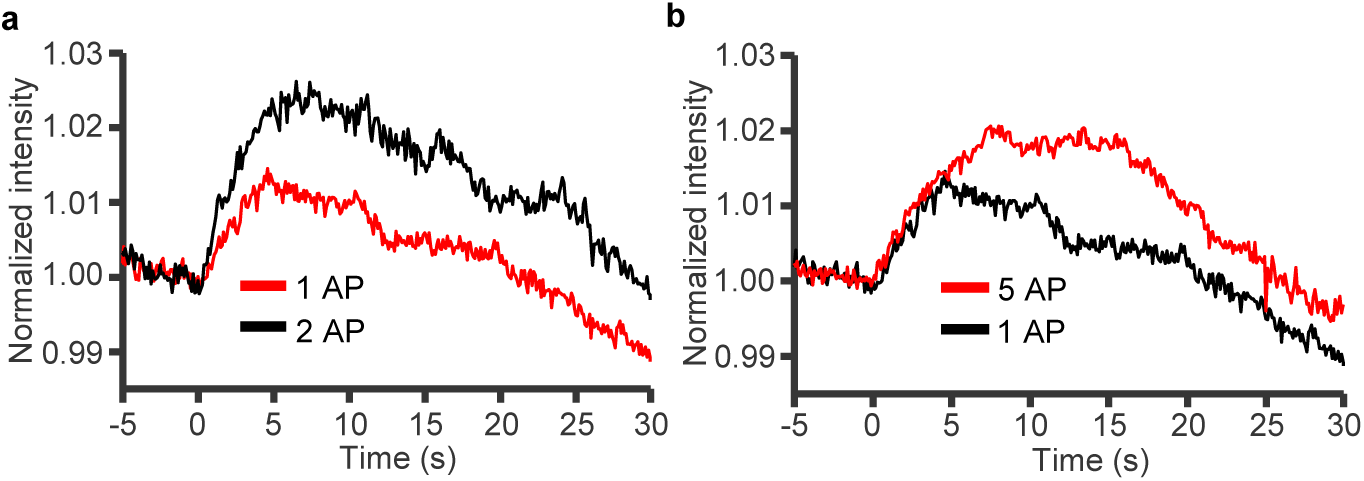
Average of individual Dynamin events in response to different stimuli. Normalized average trace of single events of endogenously tagged Dynamin I-EGFP in response to 2 AP (n = 228, black) and1 AP (n = 206, red) **(a)** and to 5 AP (n = 258, red) **(b)**. n = 4 measurement from 3 biological replicates for 1 AP and n = 6 measurements from 5 biological replicates for 2 and 5 AP.

**Extended Data Fig.6:**
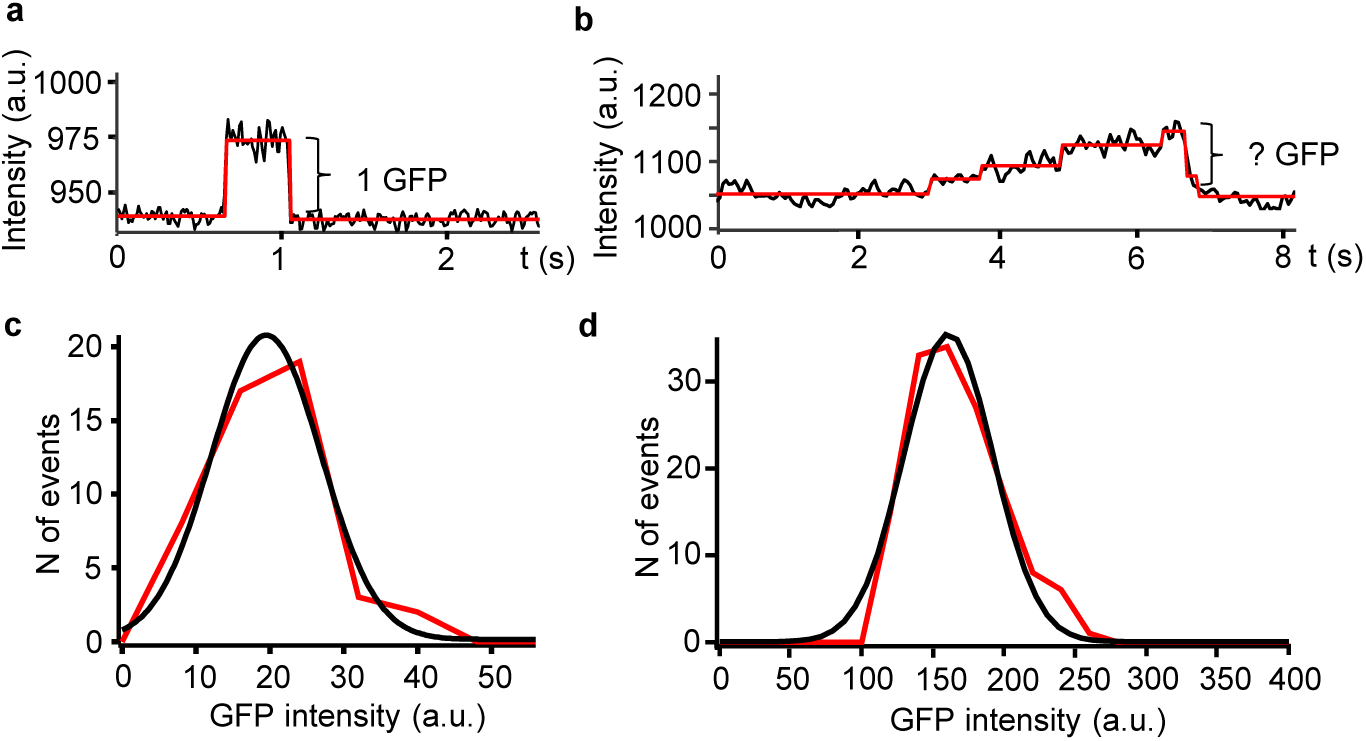
Quantification of Dynamin I molecules recruited to sites of fission. **a,** Representative fluorescence trace of a single EGFP molecule blinking event. **b,** Representative trace of evoked scission event of unknown Dynamin stoichiometry. **c,** Frequency distribution of single EGFP fluorescence amplitudes (n = 43 events). **d,** Frequency distribution of maximum fluorescence amplitudes of single Dynamin events (n = 143 events).

