## Supplementary Table 1 for "Slow scission of single synaptic vesicles by Dynamin at physiological temperature"

| | |
|---|---|
| **20bp gRNA sequences** | |
| pORANGE Dyn1 gRNA_For | CACCGCAGGTGCAGAGCGTACCGGC |
| pORANGE Dyn1 gRNA_Rev | AAACGCCGGTACGCTCTGCACCTGC |
| TKIT Dyn1 gRNA_1_For | CACCGGATGAATGAAAGGGAAGATG |
| TKIT Dyn1 gRNA_1_Rev | AAACCATCTTCCCTTTCATTCATCC |
| TKIT Dyn1 gRNA_2_For | CACCGTGAAAACTTGTGCCCCTCTG |
| TKIT Dyn1 gRNA_2_Rev | AAACCAGAGGGGCACAAGTTTTCAC |
| **PCR primers for knock-in constructs** | |
| GFPDyn1_pORANGE_PCR_For | ATAAAGCTTCCCGCCGGTACGCTCTGCACCTGGAGGTGCTAGCGTGAGCAAGGGCGAGGAG |
| GFPDyn1_pORANGE_PCR_Rev | ATACTCGAGCAGGTGCAGAGCGTACCGGCGGGCCACTTCCGTCGACCTTGTACAGCTCGTCCATGC |
| pORANGE lentiviral_For | GGACAGCAGAGATCCAGTTTGGTTAGTGAGGGCCTATTTCCCATG |
| pORANGE lentiviral_Rev | CCAGATCTGGAGCCGACACGGGTTAATGGCCATTTACCGTAAGTTATG |
| TKIT gRNA cassette_For | TAGTCTAGAGAGGGCCTATTTCCCATGATTCC |
| TKIT gRNA cassette_Rev | TCCTCGAGTCGACAATTGCT |
| **PCR primer for Halo-CLCa** | |
| Halotag_Age1_For (with Kozak seq) | CTAGACCGGTCGCCACCATGGAAATCGGTACTGGCTTTCCATTC |
| Halotag_Xho1_Rev | CTAGCTCGAGACCGGAAATTTCCAGAGTAGACAG |
